# miR-10b Mitigates Cardiac Fibrosis Associated with Aging and Myocardial Infarction via Attenuation of Lpar2 Signaling in Cardiac Fibroblasts

**DOI:** 10.64898/2026.07.02.736228

**Authors:** Evelyn-Gabriela Nastase-Rusu, Catalina-Iolanda Marinescu-Colan, Carmen Alexandra Neculachi, Ana-Mihaela Lupan, Bogdan Paul Cosman, Mihai Alin Publik, Elisa Liehn, Fabio Martelli, Mihai Bogdan Preda, Alexandrina Burlacu

## Abstract

A impairs post-infarction cardiac repair through dysregulated fibroblast activation and excessive extracellular matrix (ECM) deposition, yet the molecular mechanisms driving the age-associated defects remain poorly defined. Here, we show that miR-10b upregulation in cardiac fibroblasts acts as an endogenous cardioprotective response to myocardial infarction (MI), limiting adverse remodeling through suppression of Lpar2 (lysophosphatidic acid receptor 2). Using integrative analysis of mRNA and small RNA transcriptomes in cardiac fibroblasts from young and aged mice, we demonstrate that miR-10b is enriched in cardiac fibroblasts and further upregulated in experimental models of cardiac fibrosis, but not in hepatic fibrosis. Temporal profiling after MI revealed a biphasic regulation of miR-10b, with downregulation during the early inflammatory phase followed by upregulation during the reparative and maturation phases. Gain-of-function experiments in cardiac fibroblasts showed that miR-10b suppressed proliferation, migration, and pro-fibrotic gene expression, while promoting apoptosis under inflammatory conditions. Integrated target prediction and transcriptomic analyses identified Lpar2 as a direct miR-10b target, validated by luciferase reporter assay and confirmed at both mRNA and protein levels. miR-10b overexpression attenuated lysophosphatidic acid (LPA)-induced fibroblast proliferation and collagen I/III synthesis, supporting an anti-fibrotic role. In vivo inhibition of miR-10b in aged mice exacerbated post-infarction ventricular dilatation and wall thinning, accompanied by increased fibrotic remodeling markers, consistent with enhanced extracellular matrix remodeling, providing in vivo evidence for this regulatory axis. Collectively, these findings establish miR-10b as a protective regulator of post-infarction remodeling and in aging heart through suppression of Lpar2-mediated fibroblast activation highlighting its potential as a therapeutic target in age-associated cardiac fibrosis.

## Introduction

Cardiovascular diseases are the leading cause of death worldwide despite important advances in cardiovascular medicine. This apparent paradox is partly driven by population aging through prolonged exposure to factors with high cardiovascular risk (https://heartreport23.world-heart-federation.org/). Aging therefore represents a demographic challenge and a biological determinant of cardiovascular vulnerability ^1^. Understanding the molecular mechanisms through which aging alters cardiac homeostasis is therefore essential for identifying new strategies to prevent or limit age-associated cardiovascular disease. Among cardiovascular conditions, myocardial infarction (MI) represents one of the most common age-associated conditions. Elderly patients have an increased risk of cardiac arrest following MI and are more likely to develop mechanical complications, such as papillary muscle rupture, left ventricular free wall rupture, and ventricular septal defect ^2^. The characteristic changes observed during aging, including reduced cardiomyocyte number, impaired angiogenesis, inefficient resolution of inflammation, reduced regenerative capacity ^3, 4^ and enhanced left ventricular (LV) collagen deposition ^5, 6^ can significantly impair healing after MI, ultimately contributing to adverse cardiac remodeling and heart failure.

A key cellular player in post-MI repair process is the cardiac fibroblast. Initially considered a passive structural cell, the fibroblast is now recognized as a highly dynamic regulator of cardiac physiology and pathology, orchestrating extracellular matrix turnover, paracrine signaling, and immune responses ^7^. Its contribution becomes particularly critical after MI, when fibroblast activation and differentiation into myofibroblast lead to fibrosis and scar formation ^8^. Despite recent advances in preclinical research on post-MI remodeling, the underlying molecular pathways remain insufficiently understood ^9^. MicroRNAs (miRNAs) are important post-transcriptional regulators of gene expression and play essential roles in diverse cellular processes. Numerous studies have linked miRNA dysregulation to cardiac hypertrophy and fibrosis ^10^. However, far less is known about how aging alters miRNA expression profiles in cardiac fibroblasts and how these age-associated changes influence fibroblast behavior following MI.

Here, we investigated the functional role of miR-10b in age-associated and post-MI fibrotic remodeling. Using small RNA sequencing, in vitro functional assays, and target identification, we demonstrated that miR-10b directly targets Lpar2 and restrains fibroblast activation, proliferation, and extracellular matrix deposition. Moreover, in vivo loss-of-function approaches, we provided evidence that inhibition of miR-10b leads to an aggravated cardiac remodeling in aged mice. Together, these findings establish the miR-10b-Lpar2 interaction as a previously unrecognized regulatory mechanism in cardiac fibroblast activation and suggest that therapeutic targeting of this pathway represent a promising strategy for the treatment of age-related and injury-induced cardiac fibrosis.

## Materials and methods

### Data Availability

A detailed description of the methodology is provided in the Supplemental Material. The RNA-seq datasets are available in the Gene Expression Omnibus under accession numbers GSE279927 for long-RNA-seq and GSE153214 for small-RNA-seq. Additional data will be shared on reasonable request to the corresponding authors.

### Animals and ethics statement

All animal experiments were conducted in accordance with national and international guidelines (Directive 2010/63/EU) and approved by the National Sanitary Veterinary and Food Safety Authority according to the authorizations 389/22.03.2018, 633/21.07.2021 and 4/17.02.2023. C57BL/6J mice were purchased from The Jackson Laboratory and bred under controlled conditions, with standard housing (12 h light/dark cycle, 21 °C, 55–60% humidity) and ad libitum access to chow and water. Male and female mice were maintained under these conditions and categorized as young (2–6 months) or aged (16–25 months). A total of 309 animals were used in this study.

### Myocardial infarction

Myocardial infarction was induced in young and aged male C57BL/6 mice by permanent LCA (Left Coronary artery) ligation under ketamine/xylazine anesthesia, while sham-operated mice underwent the same procedure without ligation ^11^. Hearts were collected at defined time points after MI for molecular and histological analyses.

### Interstitial cardiac fibrosis

Interstitial cardiac fibrosis was induced by subcutaneous isoproterenol administration using either a single 100 mg/kg dose followed by tissue collection at 4 weeks or five consecutive daily injections of 200 mg/kg. Saline-treated mice served as controls, and hearts were collected for RNA extraction or histological analysis

### Hepatic fibrosis in mice

Hepatic fibrosis was induced in female C57BL/6J mice by twice-weekly intraperitoneal CCl4 injections for 14 weeks, with progressive dose escalation from 0.25 to 1.6 g/kg. One week after the final injection, liver tissue was collected for molecular and histopathological analyses.

### Echocardiography

Cardiac function and structure were assessed by transthoracic echocardiography at day 6 and day 14 after LCA ligation using a Vevo 2100 imaging system. LV volumes, wall thickness, EF, FS, SV, and CO were calculated from B-mode images using VevoLab software.

### Histology analysis

Heart and liver tissues were fixed in 4% paraformaldehyde, paraffin-embedded, sectioned at 5 μm, and stained with Picrosirius Red to assess fibrosis ^12^. Images were acquired with a Leica DMi8 microscope, and fibrotic area was quantified using LAS X software.

### Organ sample preparation

Cardiac ventricles, a portion of the right liver lobe, pancreas, left kidney, left quadriceps muscle, and whole brain from young (3 months) and old (18-25 months) mice were immediately flash-frozen in liquid nitrogen and ground to a fine powder using a mortar and pestle pre-cooled in liquid nitrogen. The resulting tissue powder was stored at −80 °C until further processing.

### Long-RNA sequencing

Long-RNA-seq was performed on total RNA extracted from cardiac fibroblasts isolated from young (4 months) and aged (16-18 months) mice after MI or sham surgery, followed by library preparation and Illumina sequencing. Differential expression analysis was performed using DESeq2.

### Small-RNA sequencing

Small-RNA-seq was performed on total RNA extracted from an independent cardiac fibroblast cohort isolated from young (4 months) and aged (16-18 months) mice, followed by small-RNA library preparation, size selection, and Illumina sequencing. Differential miRNA expression analysis was performed using edgeR.

### Cell cycle analysis

NIH/3T3 cells transfected with miR-10b mimic or scramble control were ethanol-fixed, stained with propidium iodide/RNase A, and analyzed by flow cytometry. Cell cycle distribution was determined using ModFit LT and expressed as the percentage of cells in G0/G1, S, and G2/M phases.

### Protein extraction and Western Blot

Proteins were extracted from tissue powder or cultured cells using RIPA buffer supplemented with protease/phosphatase inhibitors. Equal amounts of protein were separated by SDS-PAGE, transferred to nitrocellulose membranes, probed with specific antibodies, and quantified relative to GAPDH or total protein loading.

### Luciferase reporter assay

Wild-type or mismatched Lpar2 3′UTR fragments containing the predicted miR-10b-5p binding site were cloned into the pmirGLO dual-luciferase vector. HEK293T cells were co-transfected with reporter plasmids and miR-10b mimic, and luciferase activity was measured 24 h later.

### Isolation of cardiac myocytes and non-myocyte fractions

Adult murine cardiomyocytes were isolated from aged mouse hearts using retrograde coronary perfusion with collagenase type II, as previously described ^13^. Both cardiomyocytes and non-myocyte pellets were subsequently used for RNA isolation.

### Cardiac fibroblast isolation

Primary cardiac fibroblasts were isolated from young and aged mouse ventricles using sequential trypsin/collagenase digestion, as previously described^13^. Cells used for experimental studies were maintained between passage 0 and passage 2.

### Cell culture and transfection

NIH/3T3 cells, primary cardiac fibroblast and immortalized mouse cardiac fibroblasts were cultured under standard conditions in serum-containing medium ^13^. Cells were transiently transfected with miR-10b-5p mimic or scrambled control using Lipofectamine RNAiMAX and collected for downstream assays.

### Cell proliferation assay

Cardiac fibroblast proliferation was monitored in real time using the xCELLigence RTCA system after miR-10b or scramble transfection. Cells were treated with agoLPA2 or vehicle, and proliferation kinetics were quantified from normalized Cell Index slopes.

### Cell migration assay

Cardiac fibroblast migration was assessed using CIM-Plate 16 chambers in the xCELLigence RTCA system. Migration was recorded for 24 h and quantified from Cell Index slopes.

### Identification of miR-10b targets

The predicted target genes of miR-10b-5p were identified using four online databases: TargetScan, miRDB, DIANA (MicroT-CDS-miTG score > 7), and miRWalk. To identify potential functional targets, these genes were intersected with those significantly downregulated in the Young-MI vs Young-Sham-MI (Y-MI vs Y-Sham), Old-MI vs. Old-MI-Sham (O-MI vs. O-Sham) and Old-MI vs Young-MI (O-MI vs. Y-MI) comparisons.

### Apoptosis assay

Apoptosis was assessed in NIH/3T3 fibroblasts transfected with miR-10b mimic or scramble control using Annexin V/propidium iodide staining. Early and late apoptotic cells were quantified by flow cytometry using CytExpert software.

## Statistical analysis

Statistical analyses were performed using GraphPad Prism 9.1.2 (GraphPad Software Inc.). Two-group comparisons were analyzed using two-tailed unpaired t-tests, or one-sample t-tests when normalized values were compared with a theoretical value of 1. Multiple-group comparisons were analyzed using one-way or two-way ANOVA, as appropriate, followed by Tukey’s post hoc test unless otherwise specified in the figure legends. Repeated measurements were analyzed using two-way repeated-measures ANOVA or a mixed-effects model when values were missing. Data are presented as mean ± SD. P < 0.05 was considered statistically significant. Statistical annotations are: ns, not significant; *P < 0.05; **P < 0.01; ***P < 0.001; ****P < 0.0001.

## Results

### miR-10b is enriched in cardiac fibroblasts and upregulated with aging

We previously reported that small RNA profiling of cardiac fibroblasts isolated from young (3-mo old) and aged (18-mo old) mice revealed moderate age-dependent differences in miRNA expression with most downregulated miRNAs originating from the Meg3-Mirg locus ^13^. Subsequent analysis focused on upregulated miRNAs. Subsequent analysis was focused on upregulated miRNAs, which were conserved across vertebrates, showed expression levels above 10 reads per million (RPM), and had an adjusted p-value < 0.05. Of the 20 miRNAs meeting these criteria, five highly conserved candidates were further selected for analysis (Fig. 1A-B). Among them, miR-7a-5p and miR-29a-3p showed modest 20-30% increases with age, while miR-9-5p, miR-146b-5p, and miR-10b-5p exhibited approximately twofold upregulations (Fig. 1C). Sex-stratified analysis revealed selective upregulation of miR-10b-5p and miR-146b-5p in cardiac fibroblasts from aged males, whereas the remaining candidates showed no significant sex-dependent differences (Fig.1D).

**Figure 1.**
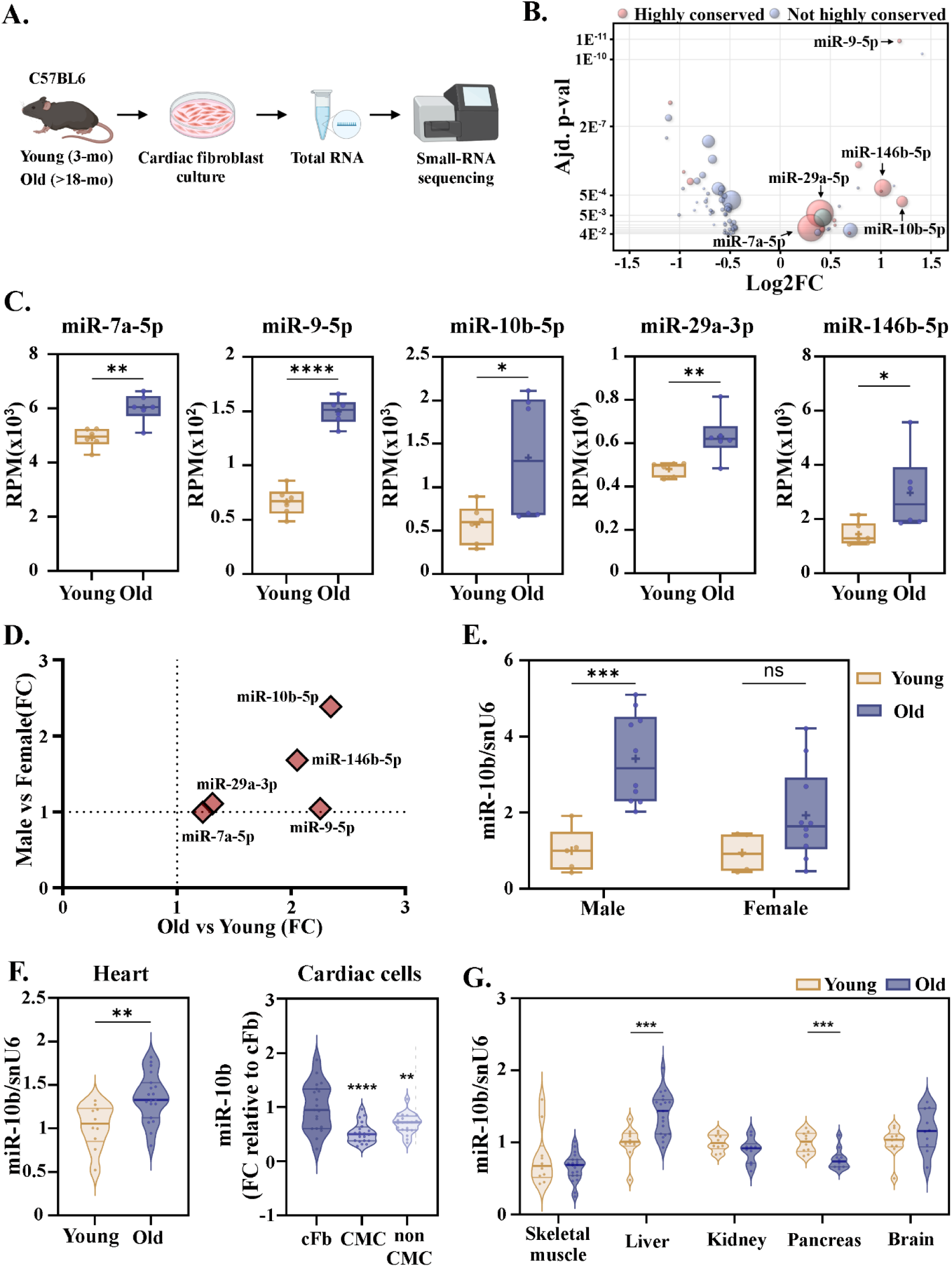
Deregulation of miR-10b during natural aging process. **(A.)** Schematic representation of experimental design. Cardiac fibroblasts isolated from young and old mice (n=6/group; n equal between male and female); **(B.)** Vulcano bubble plot showing 71 miRNAs that have over 10 RPM expression level, p.adj <0.05 and are conserved in vertebrates. The dimension of the bubbles represents the mean RPM level and the color represent different conservation status: red= highly conserved (conserved among vertebrates); blue= not highly conserved (conserved among mammals); green= ORF; **(C.)** The expression of upregulated mature miRNAs in cardiac fibroblast; **(D.)** Scatter plot showing the sex and age dependent upregulation of the selected miRNA; **(E.)** Validation of miR-10b-5p in cardiac fibroblasts isolated from young (n=5 male; 5 female) and old (n=10 male; 10 female) according to sex and age; the expression is presented relative to the young male group. Two-way ANOVA, with Tukey’s multiple comparisons test for pairs; *** p < 0.001; **(F.)** Relative expression of miR-10b in the aging heart. One-way ANOVA for equal variances with Tukey post-hoc test for pairs; * p < 0.05, ** p < 0.01, *** p < 0.001; for heart n=10-20/group; for cardiac cells n=19-20/group; graphs show means ± SD; **(G.)** Age-dependent alteration of miR-10b in several mouse organs. For each organ, the expression is presented relative to the young group; Two-tailed two-samples T-test for equal or unequal variances; Wilcoxon two-sample test for non-normal distribution;

RT-qPCR validation using an independent cohort confirmed age-dependent upregulation of miR-10b-5p only in male-derived cardiac fibroblasts. while expression in cardiomyocytes and non-myocyte cells was not sex-dependent (Fig. 1E, Fig. S1A-B).

At the whole-heart level, age-associated miR-10b upregulation was predominantly confined to cardiac fibroblasts, with significantly lower expression in cardiomyocytes and the non-myocyte fraction (Fig. 1F; Fig. S1B). Increased miR-10b expression was also detected in the liver, but not in the other tissues examined (Fig. 1G). Collectively, these data indicate miR-10b as a cardiac fibroblast-enriched, male-specific age-responsive microRNA

### Cardiac fibrosis is associated with elevated miR-10b-5p expression

Given that miR-10b upregulation was restricted to cardiac fibroblasts and detected in the aged liver, we next investigated whether miR-10b is regulated in established models of fibrosis. Liver fibrosis was induced by CCl4 administration (Fig. 2A). Picrosirius red staining confirmed normal liver architecture in control mice, whereas CCl4 treatment resulted in four time increase in collagen deposition, predominantly in perilobular regions (Fig. 2B). In contrast, miR-10b expression was not significantly altered in this model (Fig. 2C).

**Figure 2.**
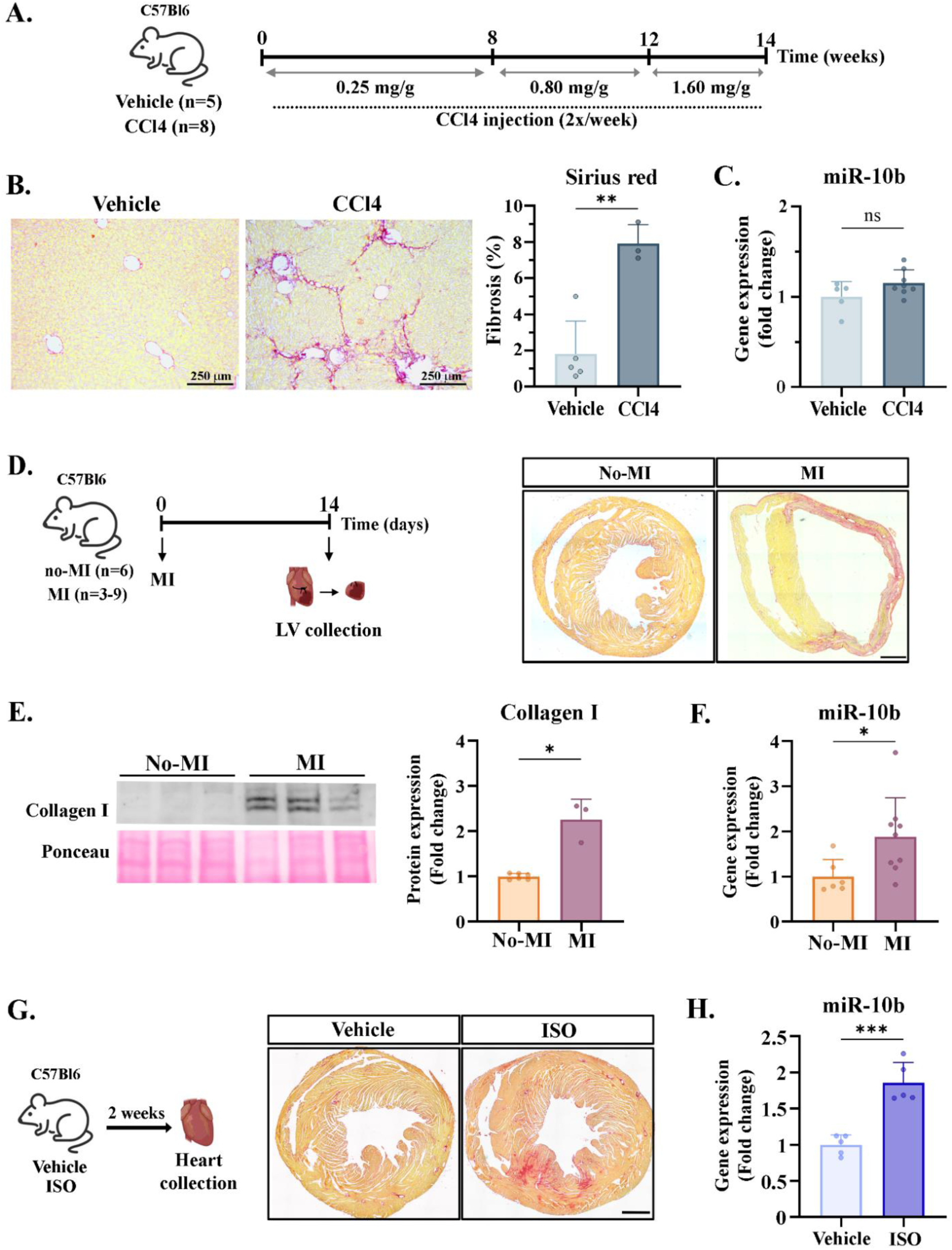
The expression of miR-10b across in vivo models of induced-fibrosis. **(A.)** Experimental design for CCl4-induced hepatic fibrosis. **(B.)** Representative Picrosirius red staining of liver tissue (left). Note the fibrotic lesion after CCl4 treatment. Relative quantification of total collagen accumulation compared to the control group (right); **(C.)** Relative quantification by RT-qPCR of miR-10b in liver sample from CCl4-challenged mice. **(D.)** Schematic representation the surgically induced fibrosis, through LCA ligation (left). Representative Picrosirius red staining of cardiac tissue after 14 days from LCA ligation (right). Note the localized fibrotic scar induced by myocardial infarction; Scale bar 1 mm. **(E.)** Representative Western blot and relative quantification of collagen type I protein levels in control and MI hearts, normalized to total protein loading. **(F.)** Relative quantification of miR-10b in MI vs no-MI samples, quantified by RT-PCR and normalized to U6 snRNA. **(G.)** Experimental design for ISO-induced fibrosis (left). Representative Picrosirius red staining of cardiac tissue after ISO administration (right). Note the diffuse fibrotic scar induced by ISO; Scale bar 0.75 mm; **(h.)** Relative quantification of miR-10b in hearts from mice receiving ISO or vehicle. Two-tailed two-samples T-test for equal variances or Welch‘s correction for unequal variances were used; graphs show means ± SD.

We next evaluated miR-10b expression in two distinct models of cardiac fibrosis. In MI model, histological examination at 14 days post-MI revealed scar tissue formation characterized by increased collagen deposition and elevated Collagen I expression, accompanied by increased miR-10b levels (Fig. 2D-E). Consistent with these findings, miR-10b expression was also increased in the isoproterenol-induced model of diffuse cardiac fibrosis (Fig. 2G-H). Collectively, these findings indicate that miR-10b is consistently upregulated in experimental models of cardiac fibrosis, irrespective of the underlying pathological mechanism, supporting a context-specific role in fibroblast-mediated fibrotic remodeling.

### miR-10b has an antifibrotic role in cardiac fibroblast

Following injury, cardiac fibroblasts become activated, proliferate, and migrate to the site of injury, where they deposit ECM that is subsequently remodeled into scar tissue during post-infarct healing ^14^. To better understand the role of miR-10b in fibrosis, we performed gain-of-function experiments in cardiac fibroblasts, overexpressing miR-10b and assessed its effects on proliferation, migration, and ECM-related gene expression (Fig. S2). Cell cycle analysis revealed that miR-10b overexpression impaired cell cycle progression, resulting in a significant accumulation of cells in S phase and a reduction in the G0/G1 population (Fig. 3A). miR-10b overexpression significantly reduced fibroblast proliferative activity, as indicated by slower growth kinetics relative to scramble controls (Fig. 3B). Consistently, migration assays further demonstrated a modest but reproducible decrease in migratory capacity (Fig. 3C).

**Figure 3.**
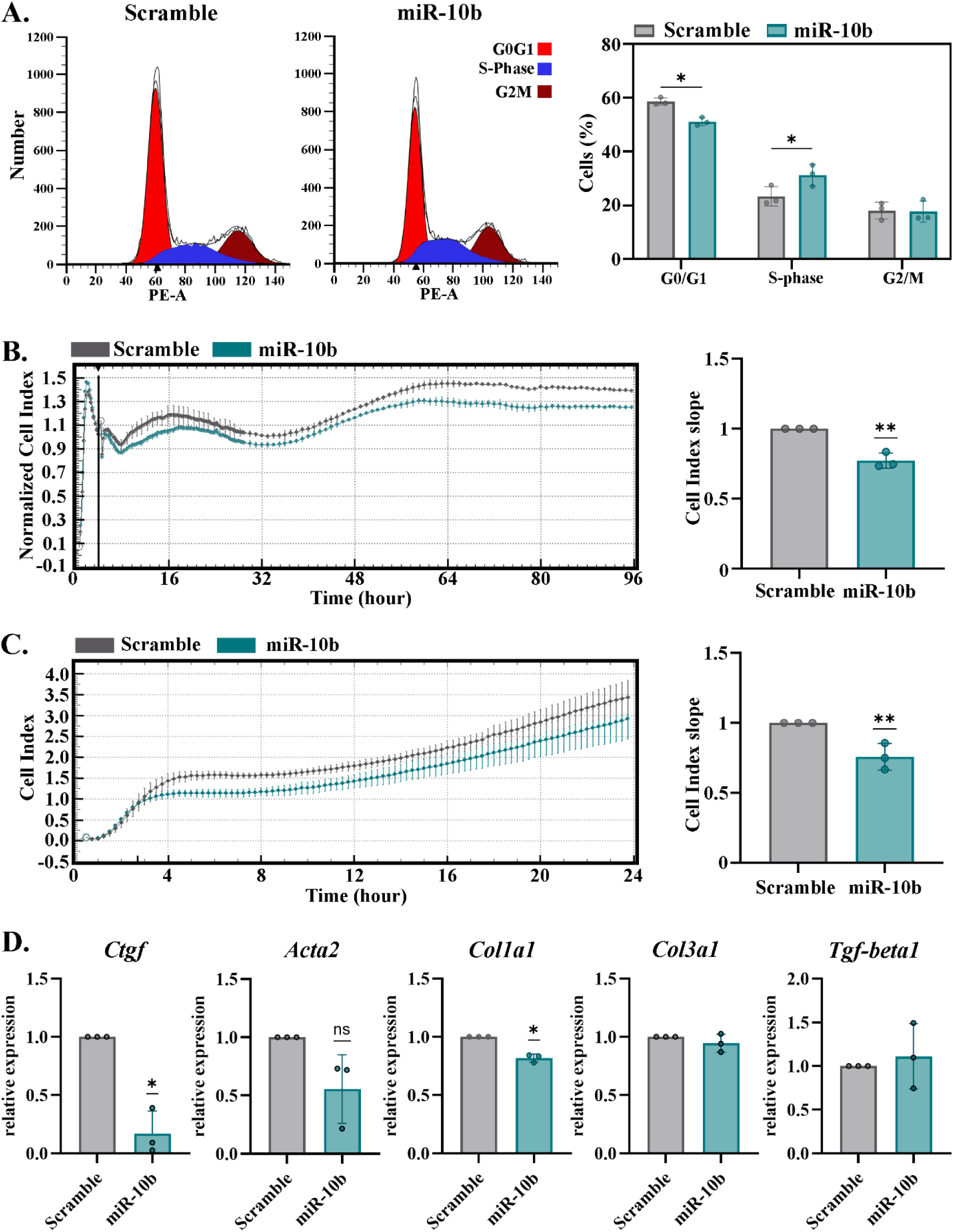
The effect of overexpressing miR-10b in fibroblasts. **(A.)** Cell cycle analyses in 3T3 fibroblast cell line overexpressing miR-10b. Histogram plots illustrate relative fluorescence intensity analyzed using ModFit LT software from one representative experiment. Bar plot shows the mean percentage of cells in G0/G1, S, and G2/M phases (n= 3 independent experiments). Two-way ANOVA, with Šídák’s multiple comparisons test for pairs. (**B-C.)** Analysis of cardiac fibroblasts overexpressing miR-10b using xCELLigence RTCA Biosensor technology. Representative Cell Index curves are shown as mean ± SD from 2–3 technical replicates per condition (left); normalized slope analysis from three independent experiments (mean ±SD) obtained with cells from different aged animals (right). **(D.)** Gene expression analyses of pro-fibrotic genes after miR-10b overexpression in immortalized old cardiac fibroblasts. Each experiment was normalized to scramble control. Statistical significance was determined using One-Sample T-test.

Given the central role of cardiac fibroblasts in ECM production, we next assessed the expression of fibrosis-related genes, including *Ctgf*, *Acta2*, *Col1a1*, *Col3a1*, and *TGF-beta1*, in immortalized old cardiac fibroblasts. Notably, miR-10b overexpression led to a marked reduction in *Ctgf and Col1a1* expression (Fig. 3D). Collectively, these data indicate that miR-10b-5p attenuates cardiac fibroblast activation by limiting proliferation, migration and pro-fibrotic gene expression.

### miR-10b-5p levels exhibit a biphasic regulation during LCA-induced cardiac fibrosis

Based on our findings that miR-10b modulates the profibrotic phenotype of cardiac fibroblasts, we next examined its temporal expression pattern during post-infarction healing, assessed at 1, 3, 7 and 14 days after coronary occlusion (Fig. 4A). miR-10b levels showed a biphasic pattern following MI, with downregulation during the inflammatory phase (days 1-3) and upregulation during the proliferative and maturation phases (days 7-14) (Fig. 4B).

**Figure 4.**
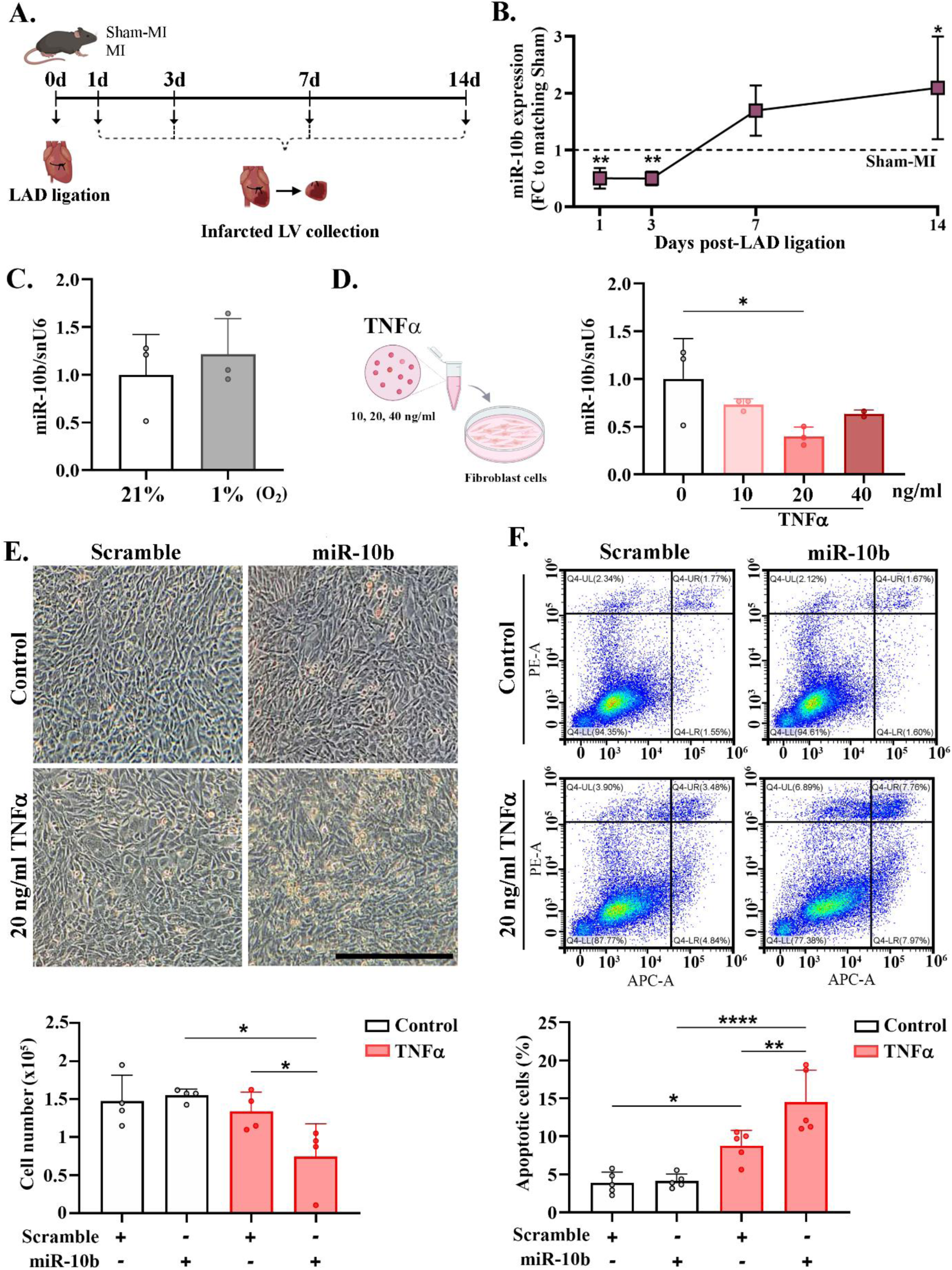
The role of miR-10b during the inflammatory phase after MI. **(A.)** Schematic representation of the experimental design. **(B.)** Relative miR-10b expression in LV. Data are calculated and compared relative to Sham samples of each time point (mean ± SD; n = 3–9; Unpaired T-test for equal variances and Welch‘s t test for unequal variances). **(C)** Relative miR-10b expression in 3T3 fibroblasts exposed to hypoxia (mean ± SD; n = 3; Unpaired T-test for equal variances). **(D.)** Schematic representation of the *in vitro* experimental design (left) and relative miR-10b expression in 3T3 fibroblasts exposed to TNF-alpha. Data are calculated and compared relative to scramble control (mean ± SD; n = 3 independent experiments; One-way ANOVA for equal variances with Dunnett’s post-hoc test for pairs); (**E.**) Representative phase-contrast images (up) and cell quantification illustrating the mortality associated to TNF-alpha exposure of miR-10b-overexpressing fibroblasts (n=4 independent experiments; Scale bare 250 mm; Two-way ANOVA with Holm-Šídák’s multiple comparisons test). **(F.)** Flow-cytometry analysis illustrating the apoptosis by Annexin V/PI staining in miR-10b-overexpressing fibroblasts after 24 h exposure to 20 ng/mL TNFα; total apoptosis = early + late apoptosis (n = 5 independent experiments); Two-way ANOVA with Holm-Šídák.

Given that miR-10b gain-of-function attenuated fibrotic activation, supporting that its upregulation may represent a compensatory mechanism to limit excessive fibrosis, we asked whether its early downregulation could be reproduced by disease-relevant stress signals. To this end, cardiac fibroblasts were exposed to TNFα or hypoxia (1% O2) for 24 hours, followed by assessment of miR-10b expression (Fig. 4C). The results showed that miR-10b levels remained unchanged under hypoxia (Fig. 4D) but were significantly reduced following stimulation with 20 ng/ml TNFα (Fig. 4E). Because fibroblast survival is essential during the early reparative response, we examined whether miR-10b modulates fibroblast susceptibility to TNFα-induced apoptosis. miR-10b overexpression increased cell detachment, enhanced apoptosis, and reduced total cell number under inflammatory conditions (Fig. 4F-G; Fig. S3A-B), consistent with the notion that its early downregulation following MI promotes fibroblast survival.

### Lpar2 is a target of miR-10b in cardiac fibroblasts

To further investigate the role of miR-10b in post-MI ventricular remodeling we assessed its expression in cardiac fibroblasts isolated from young and aged mice 7 days after Sham or MI surgery. The results showed that miR-10b was upregulated in MI-derived fibroblasts from both young and aged mice, with a more pronounced increase in the aged group (Fig. 5A).

**Figure 5.**
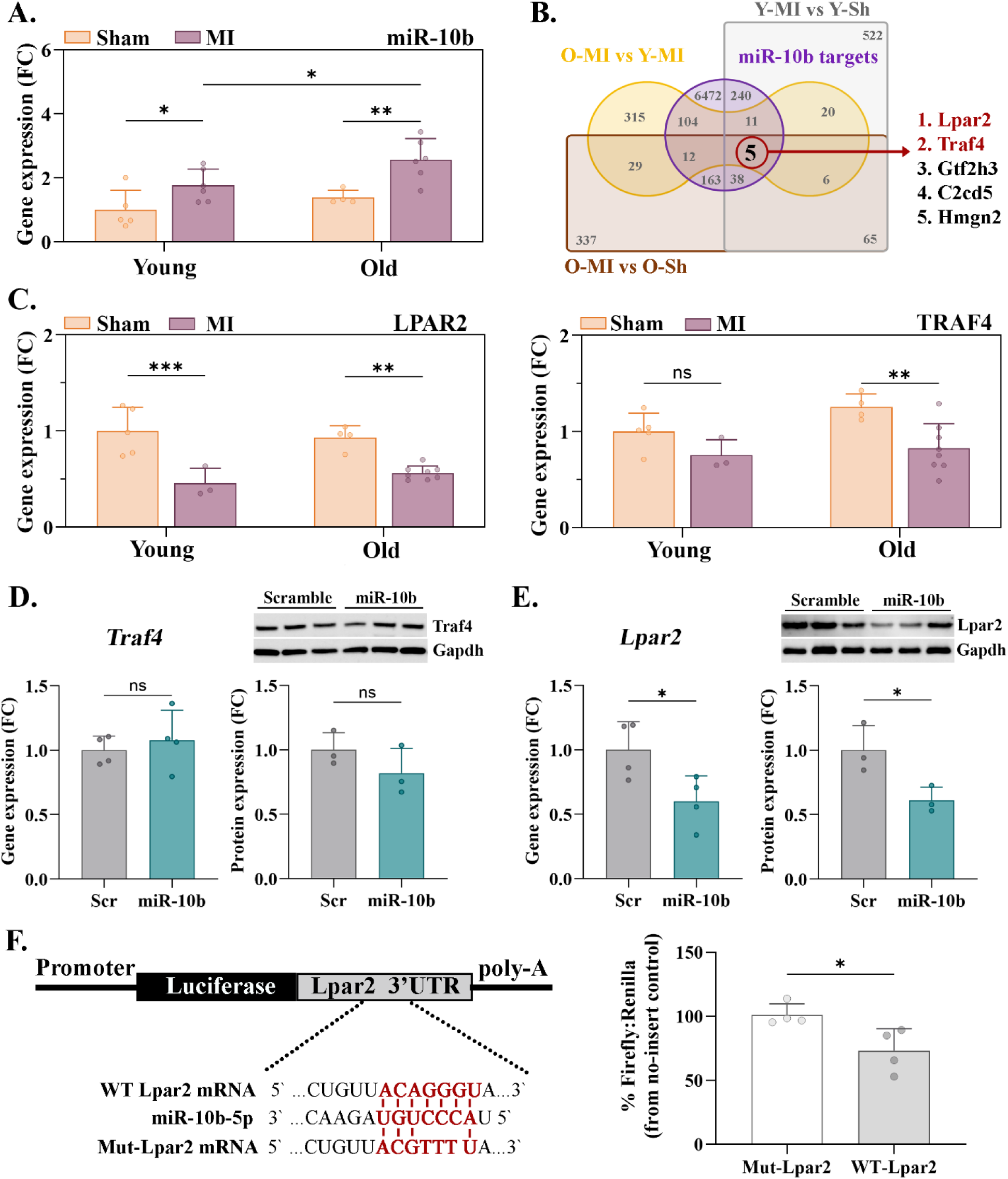
**Bioinformatic analyses and target validation of miR-10b. (A.) Relative miR-10b**in cardiac fibroblasts isolated from young and old mice 7 days post-MI; Two-way ANOVA with Fisher’s LSD. **(B.)** Venn diagram showing commonly downregulated genes of Y-MI vs Y-Sham, O-MI vs O-Sham and O-MI vs Y-MI, overlapped with miR-10b targets identified in four data bases. Fibrosis-related candidates selected for further analysis are highlighted in red. **(C.)** Relative expressions of *Lpar2* and *Traf4* in cardiac fibroblasts isolated from young and old mice at 7 days post-MI, normalized to RPL32; Two-way ANOVA with Fisher’s LSD. Protein and mRNA level of **(D.)** *Traf4* and **(E.)** *Lpar2* following miR-10b overexpression in cardiac fibroblast. Gene expression was normalized to RPL32, and protein level to GAPDH (mean ± SD; n = 3-4 independent experiments; Two-tailed two-samples T-test with Welch‘s correction for unequal variances). **(F.)** Dual-luciferase reporter. Wild-type or mismatch Lpar2-3′UTR reporter constructs were co-transfected with miR-10b mimic. Firefly luciferase activity was normalized to Renilla and expressed relative to the no-insert control (mean ± SD; n = 4 independent experiments).

Building on previous findings that aging amplifies transcriptional changes in MI-derived cardiac fibroblasts ^15^, we integrated existing mRNA sequencing data from young and aged mice after Sham or MI with predicted miR-10b-5p targets from four databases, intersecting downregulated genes across three comparisons (Y-MI vs. Y-Sham, O-MI vs. O-Sham, and O-MI vs. Y-MI). This approach identified five candidate genes, among which *Lpar2* (Lysophosphatidic acid receptor 2) and *Traf4* (TNF receptor-associated factor 4) were prioritized based on their reported involvement in fibrotic signaling pathways (Fig. 5B).

RT-qPCR validation confirmed sustained *Lpar2* downregulation in cardiac fibroblasts from both young and aged MI mice, while *Traf4* reduction was restricted to old post-MI fibroblasts (Fig. 5C). However, miR-10b overexpression reduced *Lpar2* but not *Traf4* expression (Fig. 5D-E; Fig. S4), and direct binding of miR-10b-5p to the Lpar2 3′UTR was confirmed by luciferase reporter assay (Fig. 5F).

Collectively, these findings demonstrate that miR-10b-5p directly targets *Lpar2* in cardiac fibroblasts after MI.

### MiR-10b interferes with LPA-induced cardiac fibroblast activation

Lysophosphatidic acid (LPA) is a bioactive phospholipid with growth factor-like properties that promote cell proliferation, migration, and survival. Beyond these cellular effects, LPA signaling through its receptors (LPAR1-6) has been implicated in fibrosis development ^16^.

To investigate the functional interplay between Lpar2 and miR-10b in cardiac fibroblasts, cells were transfected with miR-10b mimic and subsequently stimulated with a selective non-lipid LPAR2 agonist (agoLPA2; GRI 977143), which does not activate other LPA receptors. Live-cell analysis using the xCELLigence system showed that LPA stimulation enhanced fibroblast proliferation, as reflected by increased cell index and slope values, and this effect was abolished in miR-10b overexpressing cells (Fig. 6A). Similarly, LPAR2 activation significantly upregulated *Col1a1* and *Col3a1* expressions, whereas miR-10b overexpression attenuated this response (Fig. 6B).

**Figure 6.**
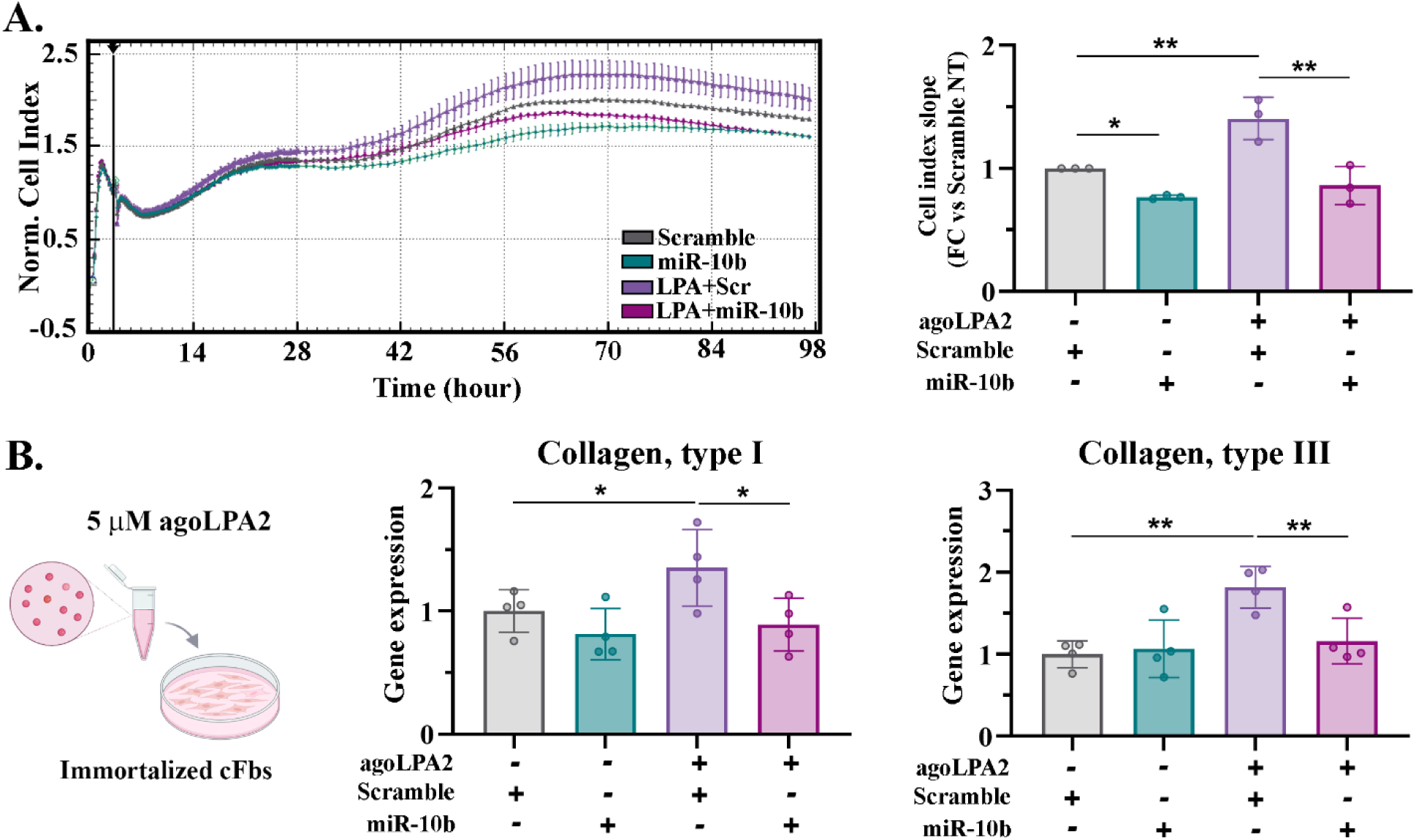
The effect of miR-10b on cardiac fibroblast proliferation and fibrotic gene expression following LPAR2 activation. **(A.)** Proliferation of aged cardiac fibroblasts analyzed by xCELLigence technology following transfection with miR-10b mimic or scramble control in the presence or absence of agoLPA2 stimulation. Representative Cell Index curves (mean ± SD of 2-3 wells) (left) and normalized slope analysis from three independent experiments (right); (mean ± SD; n = 3; Two-way ANOVA with Holm-Šídák’s multiple comparisons test). **(B.)** Experimental design and relative expression of Collagen I and III in aged immortalized cardiac fibroblasts overexpressing miR-10b after 48h of agoLPA2 exposure, normalized to S18 and expressed relative to untreated Scramble control (mean ± SD; n = 4; Two-way ANOVA with Holm-Šídák’s multiple comparisons test).

These findings further support an anti-fibrotic role for miR-10b-5p in cardiac fibroblasts that is mediated, at least in part, through *Lpar2* downregulation, thereby blunting LPA-driven signaling and limiting profibrotic activation.

### miR-10b-5p inhibition promotes adverse remodeling after MI during aging

Having identified miR-10b-5p as an age-and injury-responsive miRNA in cardiac fibroblasts, our findings support a protective role for this miRNA in limiting adverse remodeling after MI. To assess its functional relevance *in vivo*, a loss-of-function approach was employed in aged mice using a chemically modified miR-10b-specific antagomir (Anti-10b), based on the hypothesis that miR-10b inhibition would exacerbate post-MI remodeling.

Following echocardiographic confirmation of MI (Fig. S4A), aged mice received Anti-10b or scramble control at 6-and 10-days post-MI and were euthanized two weeks after surgery (Fig. 7A). Anti-10b treatment effectively suppressed miR-10b levels (Fig. 7B) and exacerbated adverse remodeling, as evidenced by increased LV area, LVID, EDV, and ESV, and reduced LVAW thickness, relative to both pre-treatment measurements and scramble controls (Fig. 7C; Fig. 4C). Although EF and FS were not significantly altered (Table 1), these structural changes suggest increased susceptibility to progressive functional deterioration. Despite the absence of significant *Lpar2* upregulation (Fig. S4B; Fig. 7D; Fig S7), miR-10b inhibition increased ECM1 and collagen levels (Fig. 7E-F; Fig. S6), consistent with the observed structural alterations, thus supporting a pro-fibrotic role for miR-10b inhibition in the aged post-MI heart.

**Figure 7.**
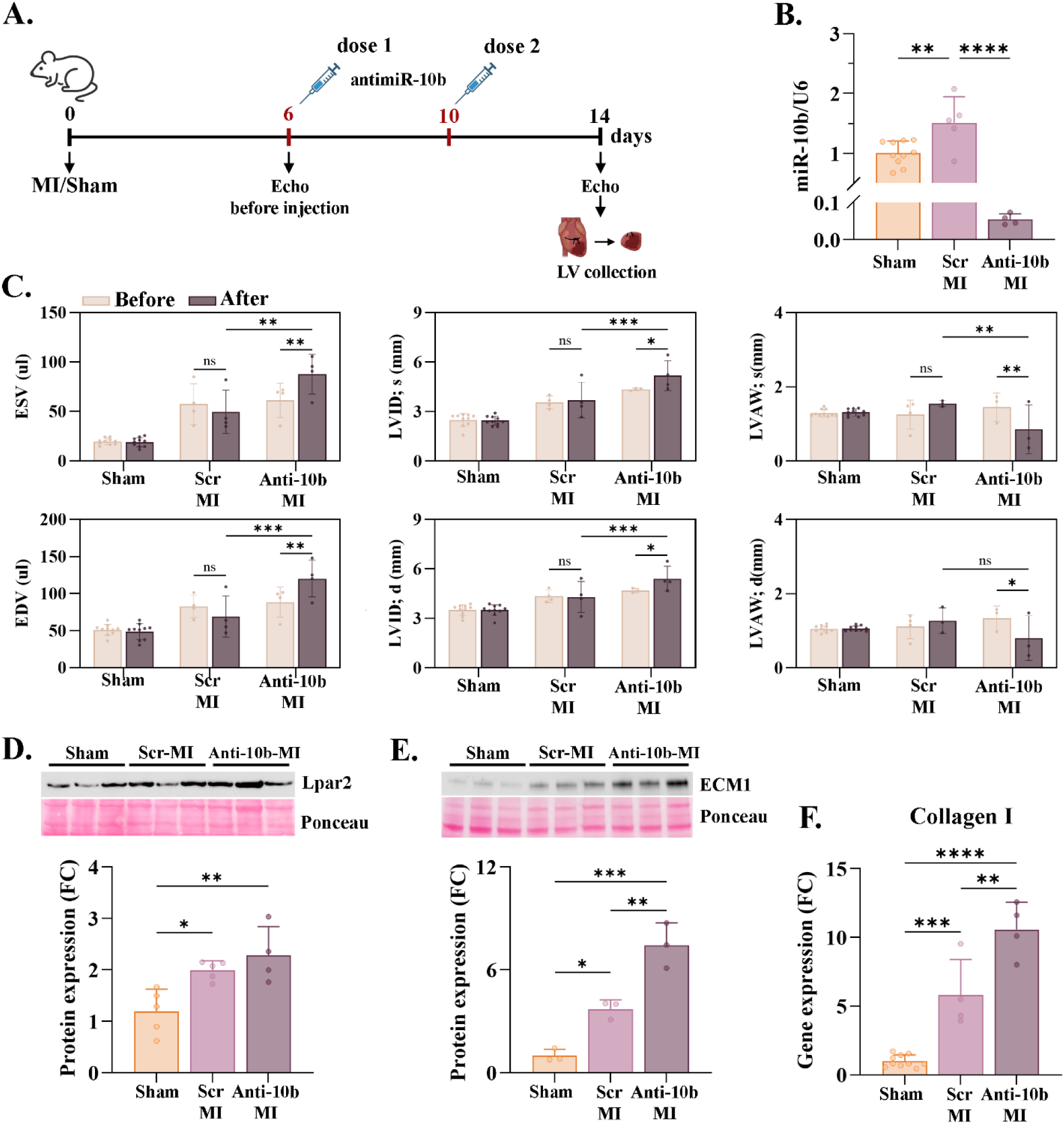
Antago-miR-10 effect of cardiac function after myocardial infraction. **(A.)** Experimental design: aged (17–18-month-old) C57BL/6 mice underwent sham or MI surgery. MI was confirmed by echocardiography 6 days post-MI, followed by administration of Scramble control (Scr) or AntagomiR-10b (Anti-10b; 10 mg/kg) on days 6 and 10 post-procedure (n = 5–10/group); **(B.)** miR-10b gene expression in LV after MI, normalized to snU6 (mean ± SD; n = 4-10; One-way ANOVA with Tukey’s multiple comparisons test); **(C.)** Echocardiographic assessment of end-systolic volume (ESV), end-diastolic volume (EDV), left ventricular internal diameter (LVID), and left ventricular anterior wall thickness (LVAW) in systole and diastole, measured before treatment initiation and at the study endpoint in sham-and MI-operated mice treated with Scr or Anti-10b. **(D.)** Representative Western blot and quantification of Lpar2 protein levels in left ventricular tissue 14d post-MI, normalized to total protein; **(E.)** Representative Western blot and quantification of ECM1 protein levels in left ventricular tissue 14 days post-MI, normalized to total protein; **(F.)** Collagen I gene expression in LV after MI, normalized to average of RPL32 and S18.

**Table 1.** Cardiac function parameters Data are presented as mean ± SD. Statistical analysis was performed using a mixed-effects model followed by Tukey’s multiple comparisons test. * vs. Before within the same group; # vs. Sham-MI at the same time point; † vs. MI-NC at the same time point. EF, ejection fraction; FS, fractional shortening; ESV, end-systolic volume; EDV, end-diastolic volume; LVAW;d, left ventricular anterior wall thickness in diastole; LVAW;s, left ventricular anterior wall thickness in systole; LVID;d, left ventricular internal diameter in diastole; LVID;s, left ventricular internal diameter in systole; LVPW;d, left ventricular posterior wall thickness in diastole; LVPW;s, left ventricular posterior wall thickness in systole; Area, left ventricular area; Area;s, left ventricular area in systole; Area;d, left ventricular area in diastole; CO, cardiac output; SV, stroke volume.

|  | Before treatment |  |  | After treatment |  |  |
| --- | --- | --- | --- | --- | --- | --- |
| Parameter | Sham | Scr-MI | anatgo-10b-MI | Sham | Scr-MI | anatgo-10b-MI |
| EF (%) | 61.83<br>±4.54 | 36.78<br>±4.91#### | 24.98<br>±9.83#####† | 60.81<br>±3.77 | 38.16<br>±8.56#### | 24.56<br>±5.38#####†† |
| FF (%) | 61.83<br>±4.54 | 36.78<br>±4.91#### | 24.98<br>±9.83#####† | 60.81<br>±3.77 | 38.16<br>±8.56#### | 24.56<br>±5.38#####†† |
| ESV (μL) | 19.64<br>±3.31 | 57.43<br>±20.70### | 61.31<br>±17.30### | 18.94<br>±4.63 | 49.66<br>±21.76## | 87.68<br>±20.17#####††** |
| EDV (μL) | 50.98<br>±7.53 | 82.48<br>±14.97## | 88.42<br>±20.34## | 48.84<br>±10.61 | 68.79<br>±27.69 | 120.12<br>±24.83#####†††** |
| LVAW;d (mm) | 1.05<br>±0.09 | 1.12<br>±0.33 | 1.34<br>±0.32 | 1.06<br>±0.07 | 1.27<br>±0.35 | 0.80<br>±0.60* |
| LVAW;s (mm) | 1.29<br>±0.10 | 1.25<br>±0.39 | 1.45<br>±0.38 | 1.31<br>±0.09 | 1.55<br>±0.08 | 0.86<br>±0.66†††** |
| LVID;d (mm) | 3.51<br>±0.32 | 4.34<br>±0.37### | 4.69<br>±0.13#### | 3.50<br>±0.33 | 4.30<br>±0.94### | 5.41<br>±0.75#####†††* |
| LVID;s (mm) | 2.50<br>±0.39 | 3.57<br>±0.38## | 4.36<br>±0.08#####† | 2.45<br>±0.27 | 3.70<br>±1.08### | 5.18<br>±0.91#####†††* |
| LVPW;d (mm) | 0.80<br>±0.09 | 0.88<br>±0.07 | 0.74<br>±0.17 | 0.84<br>±0.10 | 0.71<br>±0.19 | 0.68<br>±0.25 |
| LVPW;s (mm) | 1.10<br>±0.11 | 1.12<br>±0.17 | 0.71<br>±0.09#† | 1.12<br>±0.11 | 0.96<br>±0.39 | 0.67<br>±0.27## |
| LV Area (mm<sup>2</sup>) | 16.13<br>±5.53 | 25.04<br>±2.01# | 27.19<br>±4.47## | 17.12<br>±5.66 | 21.82<br>±4.44 | 34.92<br>±4.71#####††* |
| LV Area;d (mm<sup>2</sup>) | 21.45<br>±1.72 | 27.98<br>±3.52# | 29.39<br>±5.11## | 20.57<br>±2.79 | 24.42<br>±5.61 | 34.92<br>±4.71#####††* |
| LV Area;s (mm<sup>2</sup>) | 11.64<br>±1.35 | 21.93<br>±4.33### | 23.07<br>±5.16#### | 11.42<br>±1.87 | 20.23<br>±5.28### | 28.84<br>±3.89#####††* |
| CO (mL/min) | 18.27<br>±3.42 | 19.03<br>±1.37 | 17.74<br>±5.61 | 17.65<br>±3.15 | 20.17<br>±8.15 | 19.16<br>±1.13 |
| SV (uL) | 40.39<br>±5.15 | 37.96<br>±4.96 | 32.55<br>±10.43 | 39.97<br>±6.25 | 38.65<br>±10.12 | 41.82<br>±4.15 |

## Discussion

Non-coding RNAs play important roles in the regulation of cardiac physiological and pathological processes. However, the specific role of the vast majority of ncRNAs is still under investigation. The main findings of this study were as follows: (i) miR-10b was upregulated in the aged heart, primarily due to its increased expression and enrichment in cardiac fibroblasts; (ii) miR-10b exhibited a biphasic regulation following MI, with reduced expression during the inflammatory phase and increased expression during the reparative and maturation phases of MI; (iii) Lpar2 was identified and validated as a direct target of miR-10b; (iv) the miR-10b–Lpar2 axis restrained the profibrotic phenotype of cardiac fibroblasts; and (v) *in vivo* inhibition of miR-10b using a specific antagomir exacerbated the adverse post-MI remodeling observed in aged mice. Among the five miRNAs initially selected in this study, only miR-10b-5p was validated as being upregulated in the aged heart, primarily due to its enrichment in cardiac fibroblasts rather than cardiomyocytes. Moreover, the upregulation of miR-10b in male-derived cardiac fibroblast was particularly striking, suggesting the existence of sex-specific regulatory mechanisms that may be driven by differences in hormonal signaling, immune responses, or epigenetic regulation during aging ^17, 18^. miR-10b has been primarily studied in the context of cancer biology ^19^ and its role in cardiac fibroblast function or cardiac aging remains poorly understood. In the context of aging, dysregulated miR-10b has been associated with various neurodegenerative diseases^20^ ^21^ where it shows distinct stage-dependent patterns.

The miR-10 family, consisting of miR-10a and miR-10b, has been implicated in several fibrotic disorders, although the precise role remains incompletely understood. In this study, miR-10b-5p was upregulated in mouse cardiac tissue following MI and chronic β-adrenergic stimulation (ISO) but remained unchanged in CCl₄-induced liver fibrosis. Increased miR-10b expression has also been reported in renal ^22^, pulmonary ^23^ and oral submucous fibrosis ^24^, as well as in patients with hypertrophic cardiomyopathy ^25^, where circulating miR-10b has been proposed as a biomarker of diffuse myocardial fibrosis. Given that fibrosis in the injured heart is primarily driven by the activation of cardiac fibroblasts, these observations suggest that miR-10b may contribute to the regulation of fibroblast behavior during pathological remodeling

Cardiac fibroblast activation is essential for myocardial repair and involves a phenotypic transition toward matrix-producing myofibroblasts, characterized by enhanced proliferation, migration, and ECM secretion. In this study, miR-10b acted as a negative regulator of cardiac fibroblast activation under basal culture conditions, inducing cell cycle arrest, reducing proliferation and impairing migratory capacity. Notably, the effects of miR-10b on cell proliferation and migration have been shown to vary across different cell types and disease models. In several cancer types, including cervical, liver, gastric cancers ^26^ and myeloma ^27^, miR-10b inhibits proliferation and migration by targeting oncogenic factors such as SKA1, IGF1R, TIAM1 and MAPRE1 ^28, 29^. In contrast, in breast, bladder, and glioma, miR-10b enhances proliferation and migration through repression of tumor suppressors, including HOXD10, PTEN, and KLF4 ^30–32^, thus suggesting a tumor-supportive role. Similarly, in human embryonic stem cell–derived cardiomyocytes, miR-10b promotes proliferation via modulation of the Hippo pathway through repression of LATS1 ^33^. Together, these opposing findings point to a highly context dependent function of miR-10b, influenced by the specific cellular environment, gene expression landscape and disease state. Additionally, in our study miR-10b downregulated *Ctgf* and *Col1a1*, a key pro-fibrotic mediator. This anti-fibrotic role is supported by a previous study showing that ECFC-derived exosomes enriched in miR-10b reduced cardiac fibroblast activation, including TGF-β–induced Col-Iα and α-SMA expression ^34^.

MicroRNAs regulate multiple stages of disease progression and can show time-dependent, biphasic expression patterns that drive distinct biological responses ^35^.In line with this, in our study, miR-10b showed biphasic regulation after MI, with early downregulation during the inflammatory phase followed by upregulation during reparative remodeling. Consistently, TNFα stimulation reduced miR-10b in cardiac fibroblasts, mirroring the acute *in vivo* decrease. This inflammatory link is supported by studies showing that miR-10b-5p downregulation is associated with increased inflammation in doxorubicin-induced heart failure and allergic rhinitis, whereas its restoration attenuates inflammatory responses ^36, 37^. In our study, miR-10b overexpression under inflammatory conditions increased apoptosis and reduced cardiac fibroblast viability, suggesting that it may limit fibroblast expansion during scar maturation. In contrast, miR-10b has been reported to protect cardiomyocytes from hypoxia-induced apoptosis ^38^. Since hypoxia did not alter miR-10b levels in our cardiac fibroblasts, these findings support a cell type-specific role of miR-10b in the injured heart.

In this study we demonstrate that miR-10b is upregulated in cardiac fibroblasts following MI, with a more pronounced increase in aged hearts. Integrative target prediction and transcriptomic analyses identified *Lpar2* as a putative miR-10b-regulated gene, which was subsequently validated as a direct target. To our knowledge, this is the first evidence that age-associated upregulation of miR-10b contributes to *Lpar2* downregulation in MI-derived cardiac fibroblasts. Beyond its general effects on cell proliferation, migration and survival, LPA signaling has emerged as an important regulator of fibroblast biology, influencing proliferation, migration, survival, and fibrotic remodeling across multiple tissues influencing by promoting fibroblast activation and pathological ECM accumulation ^39, 40^. Mechanistically, we identified a novel regulatory axis in which miR-10b directly targeted *Lpar2*, linking age-associated miR-10b induction to altered LPA2-mediated signaling in cardiac fibroblast after MI. These findings support a functional interaction between miR-10b and LPA-induced profibrotic signaling pathways. This is biologically relevant given that LPA signaling promotes fibroblast activation and myofibroblast differentiation through canonical pro-fibrotic pathways, including MAPK/ERK, PI3K/AKT, and TGF-β–dependent signaling ^41–43^ our findings position the miR-10b/Lpar2 axis as a previously unrecognized modulator of post-infarction fibroblast remodeling.

Our functional assay showed that, in the presence of an LPA₂ agonist, miR-10b reduced cardiac fibroblast proliferation and collagen I/III expression, supporting a functional interaction between miR-10b and Lpar2. These findings are consistent with accumulating evidence implicating the ATX–LPA signaling axis in fibroblast activation and fibrotic remodeling. In particular, LPA2 knockdown reduced LPA-induced profibrotic activation in human lung fibroblasts ^41^, whereas Lpar1 deficiency attenuated renal fibrosis following unilateral ureteral obstruction.^42^ Similarly, genetic ablation of LPA1 attenuated hypertrophy and fibrosis in a mouse model of hypertrophic cardiomyopathy ^43^

In aged mice, miR-10b inhibition exacerbated post-MI ventricular dilatation and wall thinning, supporting a protective role for endogenous miR-10b in limiting adverse remodeling. Although Lpar2 expression was not significantly altered at this time point, miR-10b inhibition increased collagen I and ECM-1 levels, consistent with enhanced fibrotic remodeling. These structural changes align with prior cardioprotective effects of miR-10b-5p upregulation ^38^ and extend its role to post-infarction remodeling in the aged heart. LPA signaling is known to promote post-MI hypertrophy, fibrosis, and adverse remodeling ^44, 45^ although its effects are receptor-and context-dependent ^46, 47^. Our finding that miR-10b inhibition upregulated Lpar2 and worsened remodeling in aged hearts suggests that, during aging, pro-fibrotic signaling in fibroblasts outweigh the pro-angiogenic effects mediated by endothelial LPAR2. Elevated post-MI LPA levels ^47^ may further amplify fibrotic remodeling through activation of multiple LPARs.

In conclusion, our study identifies miR-10b as a context-dependent regulator of post-MI cardiac remodeling, with distinct age-and cell type–specific effects. We provided evidence that, in cardiac fibroblasts, miR-10b directly targets Lpar2, restraining fibrotic activation in the aged heart, and its endogenous upregulation limits adverse structural remodeling after infarction. Collectively, these findings position miR-10b as a potential therapeutic target for attenuating maladaptive remodeling and underscore the need for age-informed, cell type–specific strategies when targeting LPA–LPAR2 axis.

## Significance

### What is Known?

- Aging promotes baseline cardiac fibrosis while impairing post-MI repair, resulting in defective scar formation and adverse ventricular remodeling.
- LPA-signaling drives cardiac fibroblast proliferation and collagen synthesis after MI, yet the specific contribution of Lpar2 to cardiac fibroblast activation and post-MI remodeling remains undefined.
- miR-10b-5p has been implicated in cardiomyocyte responses to cardiac injury, yet its role in cardiac fibroblasts and age-associated post-infarction remodeling remains unknown.

### What New Information Does This Article Contribute?

- miR-10b is enriched in cardiac fibroblasts and exhibits biphasic regulation following MI, with early downregulation during the inflammatory phase followed by upregulation during the reparative phase.
- miR-10b suppresses cardiac fibroblast proliferation and collagen synthesis by directly targeting Lpar2, identifying the miR-10b–Lpar2 interaction as a previously unrecognized regulator of cardiac fibroblast activation.
- In aged mice, miR-10b inhibition exacerbates post-infarction ventricular dilatation, wall thinning, and adverse cardiac remodeling, demonstrating a protective role for miR-10b in the aging heart.

Aging impairs post-infarction cardiac repair through poorly understood mechanisms. Using transcriptomic profiling, gain-of-function studies, and in vivo inhibition in aged mice, we identify miR-10b as an endogenous regulator of cardiac fibroblast activation and post-infarction remodeling. miR-10b is enriched in cardiac fibroblasts and shows biphasic regulation after MI, with early downregulation during inflammation followed by upregulation during repair. Mechanistically, miR-10b restrains fibroblast proliferation and collagen synthesis by directly targeting Lpar2, thereby limiting LPA-driven profibrotic signaling. In aged mice, miR-10b inhibition worsens ventricular dilation, wall thinning, and adverse remodeling, supporting a protective role during cardiac repair. These findings identify the miR-10b-Lpar2 axis as a regulator of fibroblast activation and a potential target for limiting age-related and injury-induced cardiac fibrosis.

## Acknowledgements

This work was supported by Romania’s National Recovery and Resilience Plan (PNRR-III-C9–2022-I8, CF 186/24.11.2022-contract 760062 / 23.05.2023) and Romanian National Authority for Scientific Research and Innovation, CCCDI – UEFISCDI (COFUND-ERA-CVD-EXPERT-65/2017 and PN-III-P4-ID-PCE-2020-1340). F.M. was also partially supported by Ricerca Corrente funding from Italian Ministry of Health to IRCCS Policlinico San Donato (#1.07.128; #1.07.127), the Italian Ministry of Health (POS-T4 CAL.HUB.RIA T4-AN-09), the European Union (Next Generation EU-NRRP M6C2 Inv. 2.1 PNRR-MAD 2022-12375790 and PNRR-MCNT2-2023-12377983), and Fondazione Malattie Miotoniche ETS-Fondo Monica Stupino. The authors acknowledge the use of Claude (Anthropic, accessed 2025) for assistance with literature search support and writing refinement. All AI-assisted content was critically reviewed, verified, and revised by the authors, who take full responsibility for the accuracy and integrity of the manuscript.

## Author Contributions

EGRN, BMP, CIM, ML, and CAN performed the experiments. BPC and MAP performed histological staining and imaging. EL and AB contributed to the establishment of the MI mouse model and performed MI induction surgeries. EGRN, FM and AB designed the study, analysed and interpreted the data, and wrote the manuscript FM and AB acquired funding acquired funding. All authors contributed to the writing of the manuscript. AB supervised the study and approved the final approval of the manuscript.

## Data Availability Statement

The data that support the findings of this study are available from the corresponding author upon reasonable request

## References

1. Zhang Y, Gu Z, Xu Y, He M, Gerber BS, Wang Z, et al. Global scientific trends in healthy aging in the early 21st century: A data-driven scientometric and visualized analysis. Heliyon. 2024;10:e23405

2. Shih H, Lee B, Lee RJ, Boyle AJ. The aging heart and post-infarction left ventricular remodeling. J Am Coll Cardiol. 2011;57:9–17

3. Olivetti G, Melissari M, Capasso JM, Anversa P. Cardiomyopathy of the aging human heart. Myocyte loss and reactive cellular hypertrophy. Circ Res. 1991;68:1560–1568

4. Dai DF, Chen T, Johnson SC, Szeto H, Rabinovitch PS. Cardiac aging: From molecular mechanisms to significance in human health and disease. Antioxid Redox Signal. 2012;16:1492–1526

5. Mendes AB, Ferro M, Rodrigues B, Souza MR, Araujo RC, Souza RR. Quantification of left ventricular myocardial collagen system in children, young adults, and the elderly. Medicina (B Aires). 2012;72:216–220

6. Fajemiroye JO, da Cunha LC, Saavedra-Rodriguez R, Rodrigues KL, Naves LM, Mourao AA, et al. Aging-induced biological changes and cardiovascular diseases. Biomed Res Int. 2018;2018:7156435

7. Tallquist MD, Molkentin JD. Redefining the identity of cardiac fibroblasts. Nat Rev Cardiol. 2017;14:484–491

8. Talman V, Ruskoaho H. Cardiac fibrosis in myocardial infarction-from repair and remodeling to regeneration. Cell Tissue Res. 2016;365:563–581

9. Travers JG, Kamal FA, Robbins J, Yutzey KE, Blaxall BC. Cardiac fibrosis: The fibroblast awakens. Circ Res. 2016;118:1021–1040

10. Small EM, Olson EN. Pervasive roles of micrornas in cardiovascular biology. Nature. 2011;469:336–342

11. Preda MB, Burlacu A. Electrocardiography as a tool for validating myocardial ischemia-reperfusion procedures in mice. Comp Med. 2010;60:443–447

12. Suvarna KS, Layton C, Bancroft JD. Bancroft’s theory and practice of histological techniques. 2019:1 online resource (573 pages)

13. Lupan AM, Rusu EG, Preda MB, Marinescu CI, Ivan C, Burlacu A. Mirnas generated from meg3-mirg locus are downregulated during aging. Aging (Albany NY). 2021;13:15875–15897

14. Younesi FS, Miller AE, Barker TH, Rossi FMV, Hinz B. Fibroblast and myofibroblast activation in normal tissue repair and fibrosis. Nat Rev Mol Cell Biol. 2024;25:617–638

15. Marinescu-Colan CI, Nastase-Rusu EG, Neculachi CA, Martelli F, Cherry L, Preda MB, et al. From cancer to heart fibrosis - glipr1 highlights a subset of myofibroblasts responsive to mesenchymal stem cell therapy after myocardial infarction. Biomed Pharmacother. 2025;187:118087

16. Rancoule C, Pradere JP, Gonzalez J, Klein J, Valet P, Bascands JL, et al. Lysophosphatidic acid-1-receptor targeting agents for fibrosis. Expert Opin Investig Drugs. 2011;20:657–667

17. Regitz-Zagrosek V, Kararigas G. Mechanistic pathways of sex differences in cardiovascular disease. Physiol Rev. 2017;97:1–37

18. Lauria F, Iacomino G, Russo P, Venezia A, Marena P, Ahrens W, et al. Circulating mirnas are associated with inflammation biomarkers in children with overweight and obesity: Results of the i.Family study. Genes (Basel). 2022;13

19. Singh R, Ha SE, Yu TY, Ro S. Dual roles of mir-10a-5p and mir-10b-5p as tumor suppressors and oncogenes in diverse cancers. Int J Mol Sci. 2025;26

20. Hoss AG, Labadorf A, Beach TG, Latourelle JC, Myers RH. Microrna profiles in parkinson’s disease prefrontal cortex. Front Aging Neurosci. 2016;8:36

21. Lau P, Bossers K, Janky R, Salta E, Frigerio CS, Barbash S, et al. Alteration of the microrna network during the progression of alzheimer’s disease. EMBO Mol Med. 2013;5:1613–1634

22. Shuai Y, Xu N, Zhao C, Yang F, Ning Z, Li G. Microrna-10 family promotes renal fibrosis through the vash-1/smad3 pathway. Int J Mol Sci. 2024;25

23. Guiot J, Andre B, Potjewijd J, Jacquerie P, Cremers S, Henket M, et al. Association of fibrotic-related extracellular vesicle micrornas with lung involvement in systemic sclerosis. Eur Respir J. 2025;65

24. Jishnu PV, Shenoy SU, Sharma M, Chopra A, Radhakrishnan R. Comprehensive analysis of micrornas and their target genes in oral submucous fibrosis. Oral Dis. 2023;29:1894–1904

25. Fang L, Ellims AH, Moore XL, White DA, Taylor AJ, Chin-Dusting J, et al. Circulating micrornas as biomarkers for diffuse myocardial fibrosis in patients with hypertrophic cardiomyopathy. J Transl Med. 2015;13:314

26. Hu G, Shi Y, Zhao X, Gao D, Qu L, Chen L, et al. Cbfbeta/runx3-mir10b-tiam1 molecular axis inhibits proliferation, migration, and invasion of gastric cancer cells. Int J Clin Exp Pathol. 2019;12:3185–3196

27. Yoshida K, Yokoi A, Kitagawa M, Sugiyama M, Yamamoto T, Nakayama J, et al. Downregulation of mir-10b-5p facilitates the proliferation of uterine leiomyosarcoma cells: A microrna sequencing-based approach. Oncol Rep. 2023;49

28. Shen D, Zhao HY, Gu AD, Wu YW, Weng YH, Li SJ, et al. Mirna-10a-5p inhibits cell metastasis in hepatocellular carcinoma via targeting ska1. Kaohsiung J Med Sci. 2021;37:784–794

29. Hou R, Wang D, Lu J. Microrna-10b inhibits proliferation, migration and invasion in cervical cancer cells via direct targeting of insulin-like growth factor-1 receptor. Oncol Lett. 2017;13:5009–5015

30. Sun L, Yan W, Wang Y, Sun G, Luo H, Zhang J, et al. Microrna-10b induces glioma cell invasion by modulating mmp-14 and upar expression via hoxd10. Brain Res. 2011;1389:9–18

31. Xiao H, Li H, Yu G, Xiao W, Hu J, Tang K, et al. Microrna-10b promotes migration and invasion through klf4 and hoxd10 in human bladder cancer. Oncol Rep. 2014;31:1832–1838

32. Wang B, Zhang Y, Zhang H, Lin F, Tan Q, Qin Q, et al. Correction for: Long intergenic non-protein coding rna 324 prevents breast cancer progression by modulating mir-10b-5p. Aging (Albany NY). 2023;15:4560–4562

33. Xie Y, Wang Q, Gao N, Wu F, Lan F, Zhang F, et al. Mircrorna-10b promotes human embryonic stem cell-derived cardiomyocyte proliferation via novel target gene lats1. Mol Ther Nucleic Acids. 2020;19:437–445

34. Liu W, Zhang H, Mai J, Chen Z, Huang T, Wang S, et al. Distinct anti-fibrotic effects of exosomes derived from endothelial colony-forming cells cultured under normoxia and hypoxia. Med Sci Monit. 2018;24:6187–6199

35. Haramati S, Chapnik E, Sztainberg Y, Eilam R, Zwang R, Gershoni N, et al. Mirna malfunction causes spinal motor neuron disease. Proc Natl Acad Sci U S A. 2010;107:13111–13116

36. Ma Z, Lian H, Lin X, Li Y. Lncrna miat promotes allergic inflammation and symptoms by targeting mir-10b-5p in allergic rhinitis mice. Am J Rhinol Allergy. 2021;35:781–789

37. Cai H, Han Y. Silenced long non-coding rna rmst ameliorates cardiac dysfunction and inflammatory response in doxorubicin-induced heart failure in c57bl/6 mice via the modulation of the microrna-10b-5p/traf6 axis. J Physiol Biochem. 2025;81:99–110

38. Wu L, Chen Y, Chen Y, Yang W, Han Y, Lu L, et al. Effect of hif-1alpha/mir-10b-5p/pten on hypoxia-induced cardiomyocyte apoptosis. J Am Heart Assoc. 2019;8:e011948

39. Geraldo LHM, Spohr T, Amaral RFD, Fonseca A, Garcia C, Mendes FA, et al. Role of lysophosphatidic acid and its receptors in health and disease: Novel therapeutic strategies. Signal Transduct Target Ther. 2021;6:45

40. Gan L, Xue JX, Li X, Liu DS, Ge Y, Ni PY, et al. Blockade of lysophosphatidic acid receptors lpar1/3 ameliorates lung fibrosis induced by irradiation. Biochem Biophys Res Commun. 2011;409:7–13

41. Huang LS, Fu P, Patel P, Harijith A, Sun T, Zhao Y, et al. Lysophosphatidic acid receptor-2 deficiency confers protection against bleomycin-induced lung injury and fibrosis in mice. Am J Respir Cell Mol Biol. 2013;49:912–922

42. Pradere JP, Klein J, Gres S, Guigne C, Neau E, Valet P, et al. Lpa1 receptor activation promotes renal interstitial fibrosis. J Am Soc Nephrol. 2007;18:3110–3118

43. Axelsson Raja A, Wakimoto H, DeLaughter DM, Reichart D, Gorham J, Conner DA, et al. Ablation of lysophosphatidic acid receptor 1 attenuates hypertrophic cardiomyopathy in a mouse model. Proc Natl Acad Sci U S A. 2022;119:e2204174119

44. Tripathi H, Al-Darraji A, Abo-Aly M, Peng H, Shokri E, Chelvarajan L, et al. Autotaxin inhibition reduces cardiac inflammation and mitigates adverse cardiac remodeling after myocardial infarction. J Mol Cell Cardiol. 2020;149:95–114

45. Yang J, Xu J, Han X, Wang H, Zhang Y, Dong J, et al. Lysophosphatidic acid is associated with cardiac dysfunction and hypertrophy by suppressing autophagy via the lpa3/akt/mtor pathway. Front Physiol. 2018;9:1315

46. Chen H, Liu S, Liu X, Yang J, Wang F, Cong X, et al. Lysophosphatidic acid pretreatment attenuates myocardial ischemia/reperfusion injury in the immature hearts of rats. Front Physiol. 2017;8:153

47. Pei J, Cai L, Wang F, Xu C, Pei S, Guo H, et al. Lpa(2) contributes to vascular endothelium homeostasis and cardiac remodeling after myocardial infarction. Circ Res. 2022;131:388–403

